# A Cell Type Enrichment Analysis Tool for Brain DNA Methylation Data (CEAM)

**DOI:** 10.1101/2025.07.08.663671

**Authors:** Joshua Müller, Valentin Laroche, Jennifer Imm, Luke Weymouth, Joshua Harvey, Adam R. Smith, Daniel van den Hove, Katie Lunnon, Rachel Cavill, Ehsan Pishva

## Abstract

DNA methylation signatures are highly cell type-specific, yet most epigenome-wide association studies (EWAS) are performed on bulk tissue, potentially obscuring critical cell type-specific patterns. Existing computational tools for detecting cell type-specific DNAm changes are often limited by the accuracy of cell type deconvolution algorithms. Here, we introduce CEAM (Cell-type Enrichment Analysis for Methylation), a robust and interpretable framework for cell type enrichment analysis in DNA methylation data. CEAM applies over-representation analysis with cell type-specific CpG panels from Illumina EPIC arrays derived from nuclei-sorted cortical post-mortem brains from neurologically healthy aged individuals. The constructed CpG panels were systematically evaluated using both simulated datasets and published EWAS results from Alzheimer’s disease, Lewy body disease, and multiple sclerosis. CEAM demonstrated resilience to shifts in cell type composition, a common confounder in EWAS, and remained accurate across a wide range of differentially methylated positions, underscoring its flexibility. Application to existing EWAS findings generated in neurodegenerative diseases revealed enrichment patterns concordant with established disease biology, confirming CEAM’s biological relevance. The workflow is publicly available as an interactive Shiny app (https://um-dementia-systems-biology.shinyapps.io/CEAM/) enabling rapid, interpretable analysis of cell type-specific DNAm changes from bulk EWAS.

## Introduction

DNA methylation (DNAm) is fundamental to the establishment and maintenance of cell identity throughout development and homeostasis. It serves as a crucial regulatory mechanism, guiding gene expression and cellular function in a cell type-specific manner. Moreover, it has been shown that exposure to environmental stimuli such as toxins, diet, or stress can result in changes in DNAm that often vary between cell types [1].

In the context of neurodegenerative diseases (NDDs), accumulating evidence indicates that DNAm alterations play a central role in disease pathogenesis and progression, with many of these changes being confined to certain brain cell types [2-4]. For example, Shireby et al, comprehensively profiled Alzheimer’s disease (AD) brain samples along with purified cell-types by fluorescence-activated nucleus sorting (FANS), revealing that neuropathology-associated DNAm changes in AD are primarily driven by non-neuronal (glial) cell types, Similarly, cell type-specific DNAm changes have been reported in intron 1 of the *SNCA* gene in Parkinson’s Disease frontal cortex [5]. Given that most brain epigenome-wide association studies (EWAS) continue to rely on bulk tissue due to the absence of robust single-cell assays and the high costs and labor of cell sorting techniques, this highlights the need for novel analytical frameworks that account for cell types.

To overcome these challenges, two main types of computational strategies have been employed: cell type deconvolution and cell type enrichment. The primary aim of cell type deconvolution methods is to estimate the proportion of each cell type within a sample and correct EWAS results for these potential confounders. Numerous deconvolution tools have been developed and validated [6]. This correction is crucial for preventing spurious associations due to changes in cell type composition rather than true DNAm differences. However, deconvolution does not directly reveal which cell types are the source of observed methylation changes. As such, deconvolution helps to control cell type composition but offers limited interpretative value, restricting the biological context of EWAS findings.

Cell type enrichment methods, in contrast, provide direct interpretative value by comparing analysis results (e.g., EWAS hits) to annotated sets of CpGs specific to cell types. Although commonly used cell type enrichment methods exist for gene expression data, such as Expression Weighted Cell type Enrichment (EWCE) [7], built on single-cell transcriptomes of the human brain, direct adaptation to DNAm data is not feasible due to fundamental differences between these data types.

Currently, there are a few tools available that specifically assess the enrichment of DNAm signatures in particular tissues or cell types. For example, eFORGE (experimentally derived Functional element Overlap analysis of ReGions from EWAS) [8] is designed to assess the enrichment of differentially methylated positions (DMPs) in regulatory elements, such as DNase I hypersensitive sites, across a range of tissues. This approach offers valuable insight into the regulatory context of DMPs and can suggest functional enrichment. However, it does not provide true cell type-specific enrichment at the single-CpG DNAm resolution. Another tool, CellDMC, can reveal cell type-specific methylation changes by combining cell type deconvolution with reference panels of methylation profiles from purified cell types [9]. In practice, CellDMC estimates the proportions of constituent cell types in each bulk sample using reference methylomes and subsequently models DNAm changes within each cell type separately, aiming to distinguish which cell types are driving observed differences in bulk data. However, this approach is heavily dependent on the accuracy of the initial deconvolution step.

In the present study, we present a novel, validated Cell-type Enrichment Analysis for Methylation (CEAM) tool designed to identify cell-type-specific DNAm alterations from bulk brain tissue data quantified using the Illumina EPIC array. Our approach consists of three main steps: first, establishing CpG sets annotated to specific cell types, second, *in silico* validation of these CpG sets and the statistical framework using simulated DNAm data, and third, application of the tool using published NDD EWAS results. Notably, our framework is adaptable, making it suitable for application to other tissues beyond the brain. Here, we demonstrate the development, validation, and utility of our method using data from neurologically healthy elderly brain samples and NDD EWAS results, offering a user-friendly new avenue for cell type-resolved epigenetic analysis in NDD research.

## Methods

### Brain samples and preparation

Post-mortem samples of the prefrontal cortex (n = 20) and cingulate gyrus (n = 19) were obtained from neurologically healthy aged individuals form the Brains for Dementia Research (BDR) [10] and UK Brain Bank Network (UKBBN) biobanks, respectively. The mean age of participants in the prefrontal cortex cohort was 85.5 years (standard deviation (SD) = 8.2), with a male/female ratio of 8:12. For the cingulate gyrus cohort, the mean age was 79.1 years (SD = 10.7), with a male/female ratio of 10:9. Detailed methods for sample preparation, cell sorting, and DNA methylation profiling have been described previously [11]. In brief, brain tissue was processed using FANS to isolate four nuclear populations with distinct markers: neurons (NeuN+), oligodendrocytes (SOX10+), microglia (IRF8+), and other glia (NeuN– /SOX10–/IRF8–). As the other glia fraction consisted of nuclei negative for neuronal, oligodendrocyte, and microglial markers, it was considered to predominantly be enriched for astrocytic nuclei and is described as astrocytes henceforth. A detailed protocol for the nuclei purification method utilized is available at: https://www.protocols.io/view/fluorescence-activated-nuclei-sorting-fans-on-huma-36wgq4965vk5/v1.

### DNA methylation profiling

DNA isolation and bisulfite conversion were performed using the EZ DNA Methylation Direct™ Kit (Zymo Research) for each nuclei population, followed by DNAm quantification using Illumina EPIC (v1) arrays. Raw methylation IDAT files were imported into R (version 4.4.1) and preprocessed and quality control (QC) checked using the *wateRmelon* package (v2.10.0) [12]. The *bscon* function was used to exclude samples with a bisulfite conversion rate below 80%. The *outlyx* function was applied to identify and remove outlier samples, defined as those falling outside twice the interquartile range on the first principal component (PC) or having a Mahalanobis distance greater than 0.15 from the data center. The *pfilter* function was used to remove low-quality probes, based on default detection p-value and bead count thresholds. Similarly, samples with more than 1% low-quality probes were excluded. The *estimateSex* function was used to compare known sex with predicted sex. The *preprocessNoob* function was used to perform dye-bias and background correction via the Normal-exponential Out-Of-Band (NOOB) method [13]. The *BMIQ* function was applied for Beta MIxture Quantile dilation (BMIQ) normalization to harmonize the signals detected by the two different probe designs [14]. Intra-sample normalization was conducted to account for the known dye-bias of the two-color channels and the differences between type I and type II probe designs present in EPIC arrays. Probes located on sex chromosomes, cross-hybridizing probes, and probes at sites of common single nucleotide polymorphisms (SNPs) were excluded as recommended by McCartney et al. [15]. The ComBat function from the sva package (v3.52.0) [16] was used to perform batch correction on samples from the UKBBN cohort to account for differences between source Brain Banks. After pre-processing, 75 and 80 samples remained in the UKBBN and BDR cohorts, respectively, with 784,595 and 784,075 probes retained. PC analysis (PCA) was performed on centered, unscaled beta-values to assess overall data quality. PCA plots were examined to determine whether samples grouped by cell type as expected and to identify potential sources of confounding variation (**Supplementary Figure 1**).

### Establishing cell type-specific CpG sets

Cell type-specific CpG sets were constructed using DNAm data from purified cell types across two cohorts. Three levels of cell type-specificity were defined, each using progressively less stringent selection criteria:

#### High-specificity sets

CpGs were included if they exhibited intermediate median methylation (0.1 < β < 0.9) in only one of the four cell types. To ensure specificity and reduce noise, these CpGs also had to differ by at least 0.1 in median methylation from every other cell type. CpGs with intermediate level of methylation (0.1 < β < 0.9) tend to display greater cell-to-cell and inter-individual variability, making them especially relevant for detecting disease-associated changes [17]. CpGs that were consistently hypomethylated (β < 0.1) or hypermethylated (β > 0.9) across all four cell types in both cohorts were excluded.

#### Medium-specificity sets

CpGs were included if they showed intermediate methylation levels in up to three cell types, allowing annotation to multiple cell types if the criterion was met.

#### Low-specificity sets

For these sets, exclusivity was further relaxed. CpGs were included if they had intermediate methylation in a given cell type (or types) and were significantly different (P _Bonf_< 0.05) from all other cell types, as determined by generalized linear mixed models (nlme package v3.1.166), adjusting for age, sex, and individual as a random effect. For each CpG, six pairwise comparisons were performed, with Bonferroni correction for multiple testing. In addition, CpGs from the previous specificity levels were added regardless of fulfilling this specific criterion. Illustrative examples of methylation values for each level of specificity are provided in **Figure 1**.

**Figure 1.**
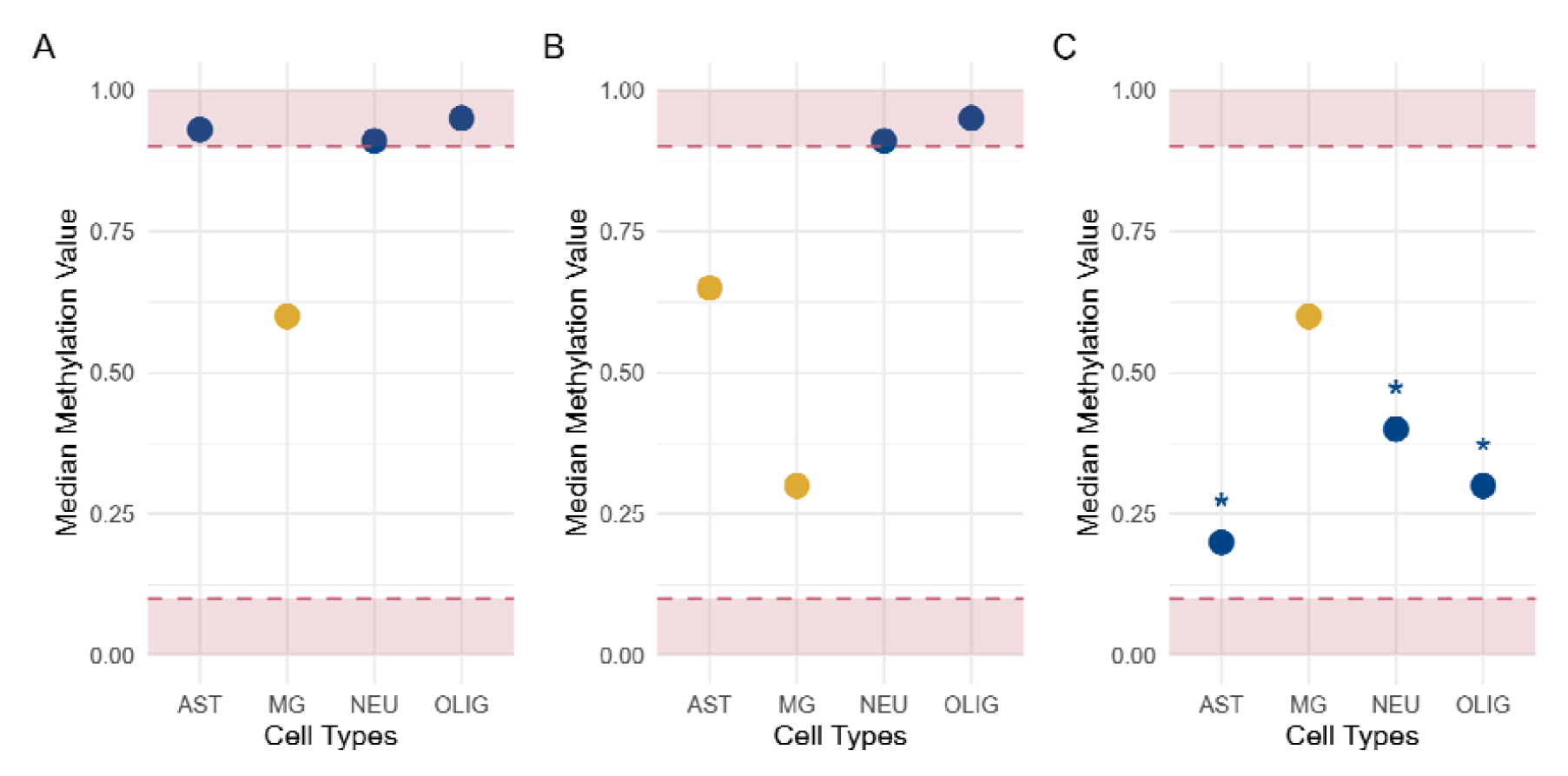
Schematic representation of methylation value criteria for CpG inclusion in target cell type-specific sets. (A) *High-specificity* CpG sets: The median methylation in the target cell type must be intermediate (outside of red areas), while all other cell types must be hypo-or hypermethylated (within red areas). (B) *Medium-specificity* CpG sets: The median methylation in the target cell type must be intermediate (outside of red areas), and at least one other cell type must be hypo-or hypermethylated (within red areas). (C) *Low-specificity* CpG sets: The median methylation in the target cell type must be intermediate (outside of red areas) and significantly different from all other cell types (indicated by *; target cell type indicated in yellow).

### *In silico* validation

#### Generating in silico DNAm datasets

To evaluate the performance of overrepresentation analysis (ORA) as the statistical method employed in the enrichment tool, we generated two identical sets of 250 “pseudo-bulk” samples, which were assigned to disease or control groups. For each sample, four cell type-specific methylation profiles were simulated using methylation patterns derived from brain purified cell DNA methylation data. Each simulated CpG was annotated as belonging either to a “set” or to the “background,” maintaining the same proportions observed in the high-confidence CpG panel derived from postmortem data in the previous step. For example, for the neuron-specific set, CpGs were simulated such that their mean methylation value in neurons was randomly drawn from a range of 0.15 to 0.85. For these same CpGs in the other cell types within the same sample, the mean values were randomly drawn from either 0.01–0.09 or 0.91–0.99, reflecting the cell-type specificity observed in high-confidence neuron-specific CpGs.

The remaining CpGs were simulated as background, with values for all cell types randomly drawn between 0.15 and 0.85 to avoid assignment to any cell type-specific set. Additionally, to mimic the heteroskedasticity typically observed in DNAm beta values, the SD of each truncated normal distribution was defined as a function of the mean.

For computational feasibility, the number of simulated CpGs was limited to 25% of a pre-processed EPIC array, resulting in 200,000 CpG sites per sample. The bulk methylation profile for each sample was then generated by combining these cell type-specific profiles.

The proportion of each cell type in each sample was randomly drawn from a Dirichlet distribution, ensuring that the weights for all cell types summed to one.

To ensure that cell type composition was the only source of variation between disease and control samples, methylation values were generated identically for both groups. However, the mixing weights used to combine the cell type-specific profiles were altered between the two groups. In the control group, the weights were drawn from a Dirichlet distribution with equal parameters for each cell type. In the disease group, the weights were increased equally for three cell types, resulting in a relative reduction of the remaining cell type in the pseudo-bulk samples.

DMPs were identified from the pseudo-bulk samples by fitting a linear regression model to each CpG site and correcting for multiple testing using the Benjamini-Hochberg method. This analysis was performed twice: once without adjusting for the known cell type proportions (the mixing weights) and once with the mixing weights included as covariates.

Lastly, to mimic the varying quality of cell type proportion estimates typically encountered in real EWAS, Gaussian noise gradually added to the mixing weights in the regression model. The resulting sets of identified DMPs were then used as input for the enrichment analysis, with odds ratios (ORs) and accompanying p-values computed using the modified Haldane-Anscombe correction as described in [18]. Simulation of DNAm data and subsequent analyses were performed over 10 iterations and multiple levels of Gaussian noise, with SD ranging from 0 to 1.5. An overview of the procedure for generating methylation values for each CpG is provided in **Supplementary Figure 2**.

#### Assessing the cell type enrichment analysis with varying input sizes

To evaluate the general sensitivity and specificity of the enrichment analysis and its applicability across a range of possible EWAS outcomes, a variable number of DMPs were introduced into the simulated DNAm data. In contrast to the previous step, for these simulations we generated 500 pseudobulk samples, divided over disease and control groups, with mixing weights sampled from a Dirichlet distribution with equal parameters. DMPs were arbitrarily assigned to CpGs within sets for one of two cell types, while the remainder were assigned to background CpGs. Specifically, half of the DMPs were assigned to the simulated neuron set, 20% to the simulated microglial set, and a minor fraction (0.1%) to the simulated oligodendrocyte and astrocyte sets. The remaining 29.8% of DMPs were distributed among background CpGs. Simulations were performed over 10 iterations, with the number of DMPs ranging from 2 up to 1,024. Only the corrected approach was applied, without the addition of noise.

#### Application for independent NDD bulk EWAS

The enrichment tool was applied to three independent bulk NDD EWAS results. First, in relation to AD, we took the 236 Bonferroni-significant CpGs that were associated with Braak neurofibrillary tangle (NFT) stage from an EWAS meta-analysis of three Illumina 450K array prefrontal cortex datasets (n=961) and tested for enrichment against 403,763 probes. Second, for Lewy Body Disease (LBD) we took the 72 CpGs that were suggestively associated (p-value < 1×10□) with Braak Lewy body (LB) stage in a recent EWAS meta-analysis of three Illumina EPIC array datasets generated in the prefrontal cortex (all three datasets) and cingulate gyrus (one dataset) (n= 1239, unique donors = 855) and tested for enrichment against all 774,310 CpGs reported in the summary statistics. Finally, to assess specificity, the approach was applied to a multiple sclerosis (MS) EWAS on white matter (n=10) [19], using 8,336 FDR significant DMPs identified between lesions and normal-appearing white matter, with 29,446 CpGs as the background. The overview of the study workflow is illustrated in **Figure 2**.

**Figure 2.**
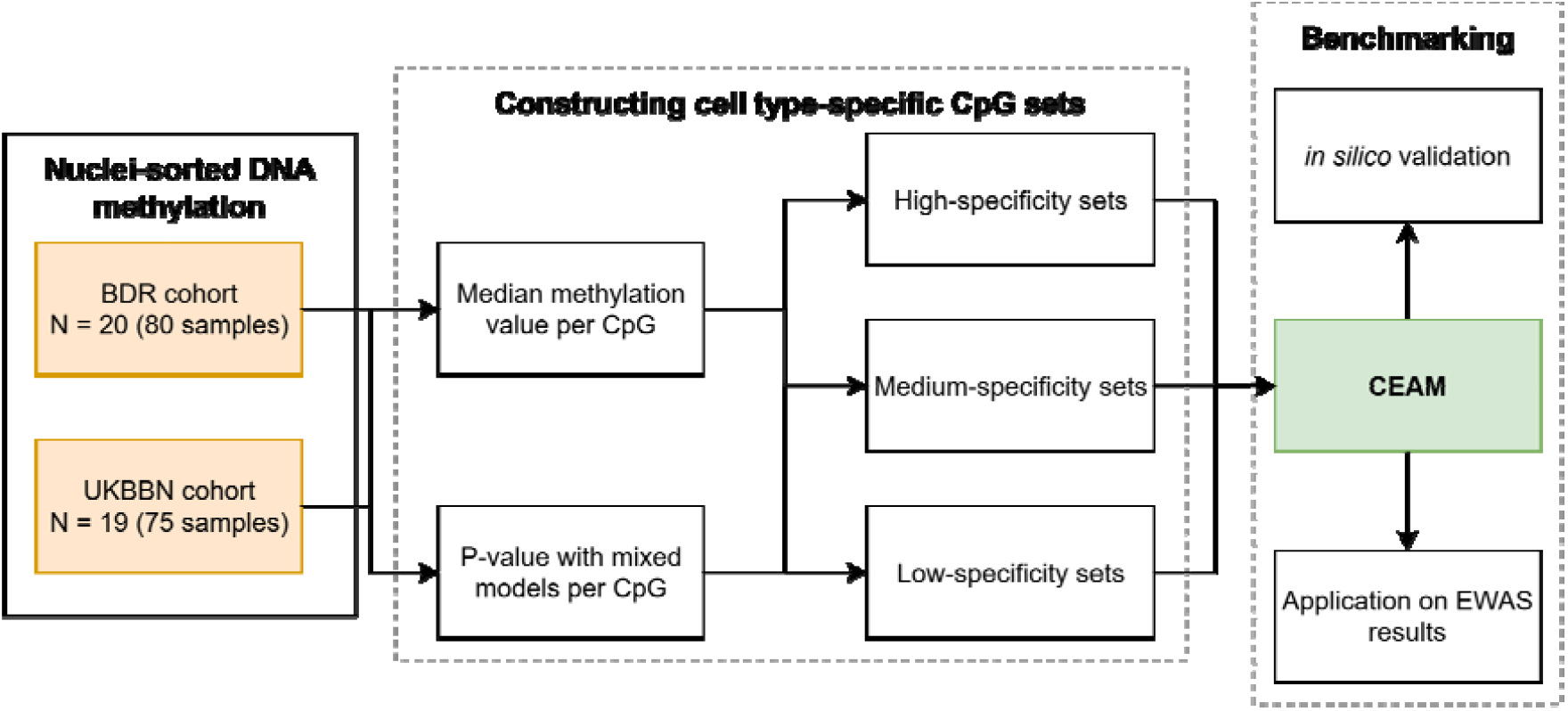
Overview of the study workflow. Nuclei-sorted DNA methylation data from post-mortem prefrontal cortex and cingulate gyrus samples were obtained from two independent cohorts of neurologically healthy aged individuals. DNA methylation profiling was performed using Illumina EPIC arrays after nuclei sorting by FANS. Following quality control and preprocessing, cell type-specific CpG sets were constructed independently in both cohorts using three levels of specificity criteria. Only CpGs present in both cohorts were retained for downstream analyses. The resulting CpG sets, together with the over-representation analysis (ORA) approach, were evaluated in two stages: (i) using *in silico* simulated pseudo-bulk datasets to assess performance under various EWAS-like scenarios and (ii) applying the method to published EWAS results from Alzheimer’s disease, Lewy body disease, and multiple sclerosis.

#### Tool Implementation and web application

The CEAM framework is implemented as a publicly available Shiny web application. All analyses described here can be reproduced via the web interface, which supports uploading user-supplied EWAS summary data and automated processing with reference panels reported in this study. The app guides users through each step and provides downloadable result files and visualizations. Source code and user documentation are available at https://github.com/Dementia-Systems-Biology/CEAM.

## Results

The high-, medium- and low-specificity CpG sets we identified differed markedly in size across cell types and specificity levels (**Figure 3**). High-specificity CpG sets for astrocytes and oligodendrocytes were at least fourfold smaller than those for neurons and microglia, consistent with greater overlap in intermediate methylation patterns between astrocytes and oligodendrocytes. In contrast, the medium- and low-specificity sets were substantially larger and exhibited more similar sizes across all cell types. These differences illustrate how the stringency of cell type specificity impacts the composition and size of CpG sets, particularly among cell types with overlapping methylation profiles. (**Figure 3, Supplementary Figures 3-4)**.

**Figure 3.**
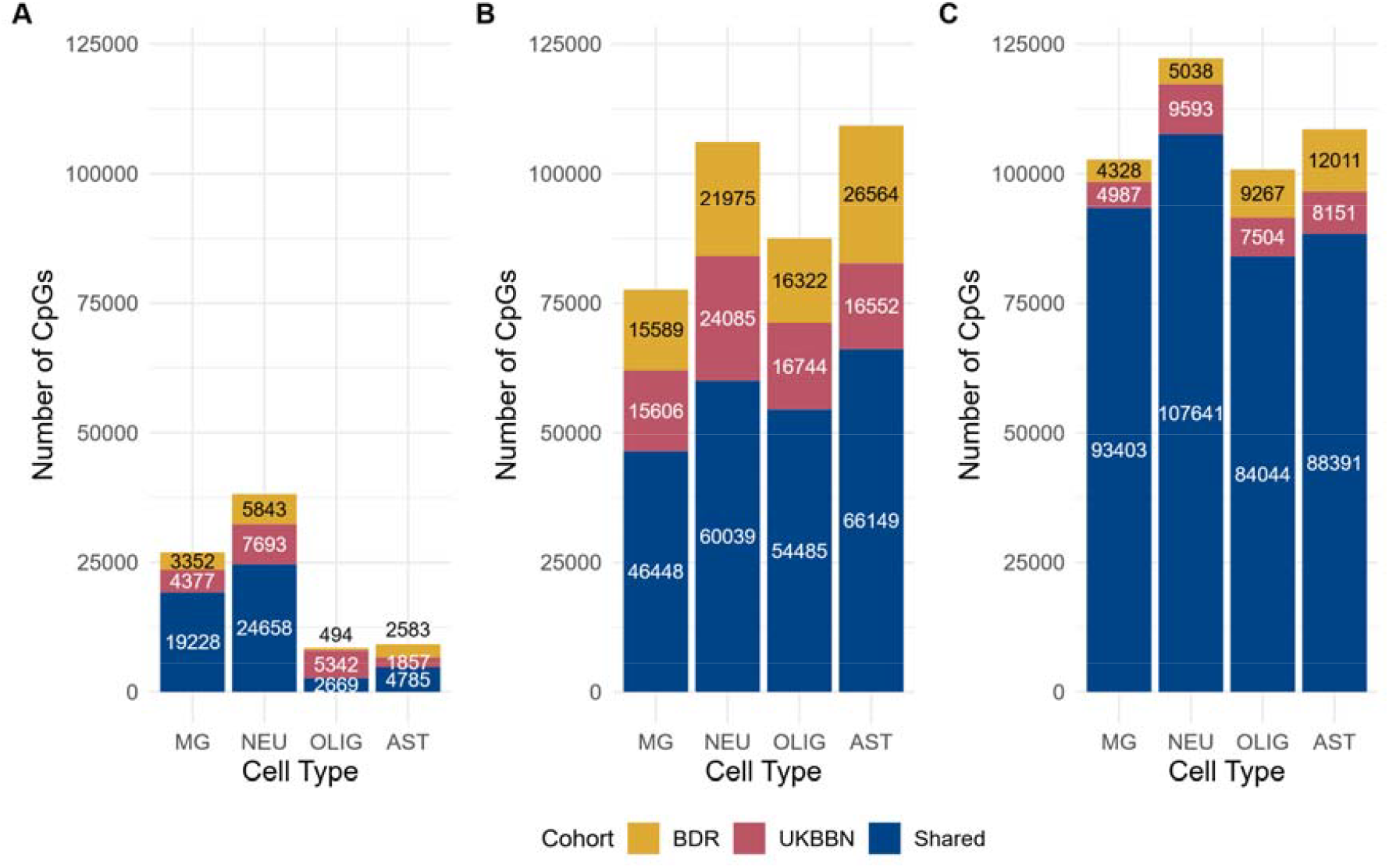
Visualization of CpG set size and composition. (A) The high-specificity CpG sets are the smallest of all, with the oligodendrocyte and astrocyte sets at least fourfold smaller than the microglia and neuron sets. (B) The medium-specificity sets increased by 2.4-fold for microglia, 2.4-fold for neurons, 20.4-fold for oligodendrocytes, and 13.8-fold for astrocytes, due to the inclusion of CpGs shared between cell types. (C) The low-specificity sets increased further in size and all cell types had comparable set sizes. All plots show the CpGs that were found in both cohorts (blue) or in only one cohort (yellow: BDR, red: UKBBN). Cell types are abbreviated as follows: MG = microglia, AST = astrocytes, NEU = neurons, and OLIG = oligodendrocytes.

The medium- and low-specificity sets were progressively larger than the high-specificity sets, as more CpGs met the less stringent criteria for inclusion. Across the entire EPIC v1 array, the high-, medium-, and low-specificity sets comprised approximately 5.9%, 13.6%, and 26.9% of all measurable CpGs, respectively. Illustrative examples of methylation distributions for each specificity level are shown in **Figure 2**. Notably, the increased inclusivity of the medium- and low-specificity sets resulted in a broader range of methylation values and greater overlap between cell types. This pattern underscores the trade-off between stringency and coverage when defining cell type-specific CpG sets.

### Shifts in cell-type composition can bias cell type enrichment results

To assess the robustness of the ORA-based enrichment tool to confounding introduced by shifts in cell type composition between disease and control groups, *in silico* DNAm datasets were generated as described in the Methods. Analyses using high-specificity CpG sets, without correction for cell type composition (i.e., omitting mixing weights as covariates), resulted in strong false-positive enrichment signals for two of the three cell types that their proportion were artificially increased in the disease group **(Figure 4)**. When the regression models were adjusted for cell type composition by including the mixing weights, this confounding was eliminated, and enrichment became non-significant for all four cell types.

**Figure 4.**
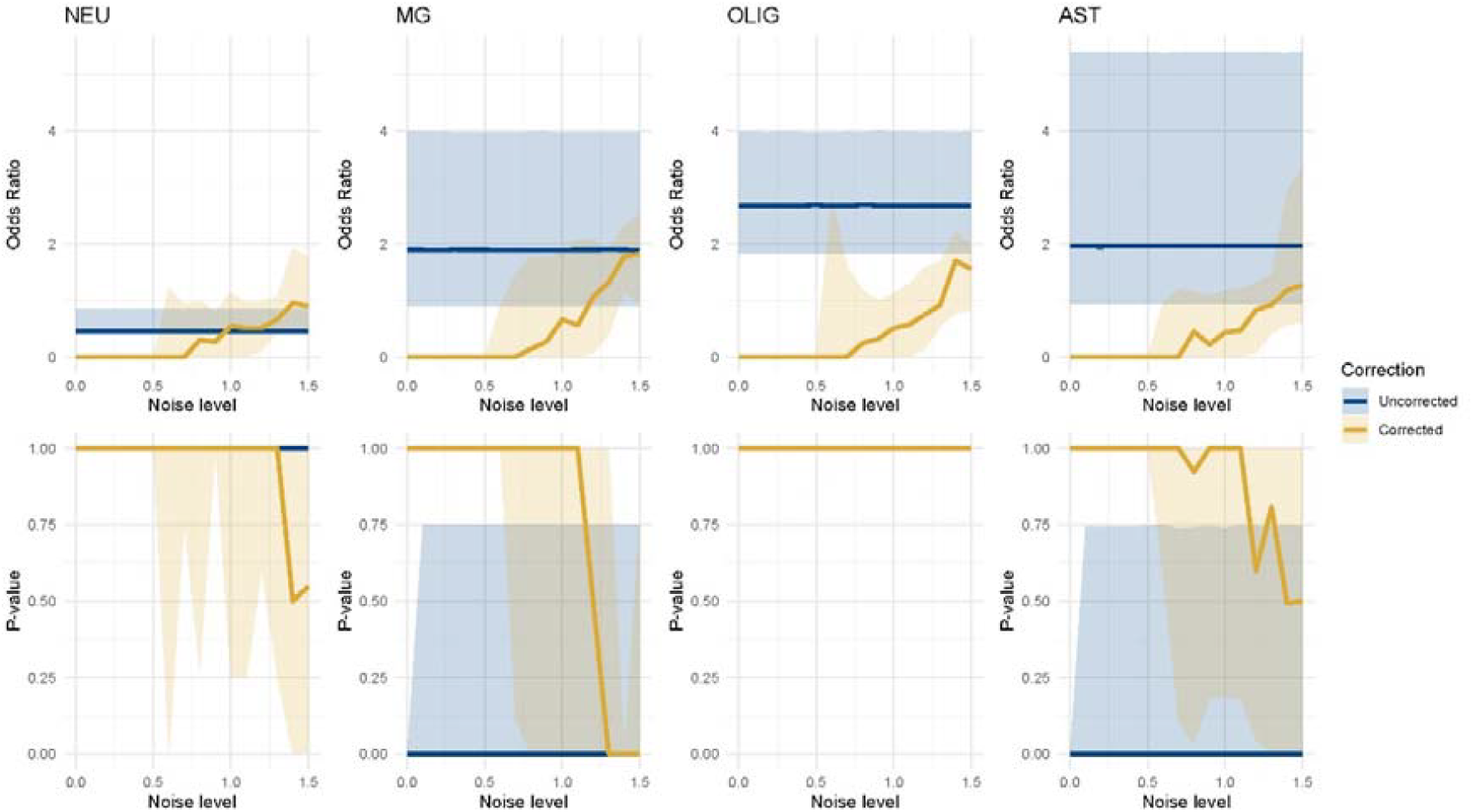
Validation on simulated data with shifted cell type proportions in simulated high-specificity CpG sets. Median odds ratio (OR) (upper panel) and p-value (lower panel) (Y-axis) over scenarios of increasing noise in cell type proportions used for correction (X-axis). Each median was computed over 10 iterations and the ribbons show the interquartile range (IQR). From left to right, the enrichment results in each simulated cell type-specific CpG set for neurons (NEU), microglia (MG), oligodendrocytes (OLIG) and astrocytes (AST), uncorrected for cell type composition (yellow) and once corrected (blue). A relative reduction of neurons in the simulated data causes microglia and astrocytes to become significantly enriched in the uncorrected scenario. However, correcting the analysis with known cell type proportions returns no enrichment in any cell type. After adding enough noise (noise level = 0.6) microglia and astrocytes become similarly enriched to the uncorrected results. In instances where not a single CpG of a set was in the input and therefore no OR could be computed, the OR was set to 0 for visualization purposes.

When Gaussian noise was added to the cell type proportion estimates used for correction, the effectiveness of this adjustment diminished. Spurious enrichment re-emerged, with a noise SD of 0.6 marking the threshold at which correction started to lose its effect. At higher noise levels, the corrected results increasingly resembled those of the uncorrected analyses, indicating that inaccurate cell type proportion estimates can substantially compromise the ability to resolve true cell type-specific enrichment patterns (**Figure 4**).

Simulations using medium-specificity CpG sets showed a similar pattern. However, the magnitude of the ORs for enrichment was considerably higher, highlighting the impact of specificity level on the sensitivity of the approach (**Supplementary Figure 5**).

### Performance across varying input sizes

To evaluate the overall performance of the enrichment analysis and its applicability to diverse EWAS outcomes, the method was applied to *in silico* DNAm datasets containing varying numbers of DMPs, ranging from 2 to 1,024. The majority of DMPs were assigned to CpGs annotated to the simulated neuron set, with a smaller fraction assigned to the microglia set, and the remainder distributed among CpGs not included in any set. The ORA approach consistently identified neurons and microglia as significantly enriched, while oligodendrocytes and astrocytes were not enriched (**Figure 5**). Remarkably, the approach was able to correctly detect the underlying enrichment patterns even with very few DMPs (fewer than 8). These results demonstrate that the ORA approach produces reliable results across a wide dynamic range of input sizes.

**Figure 5:**
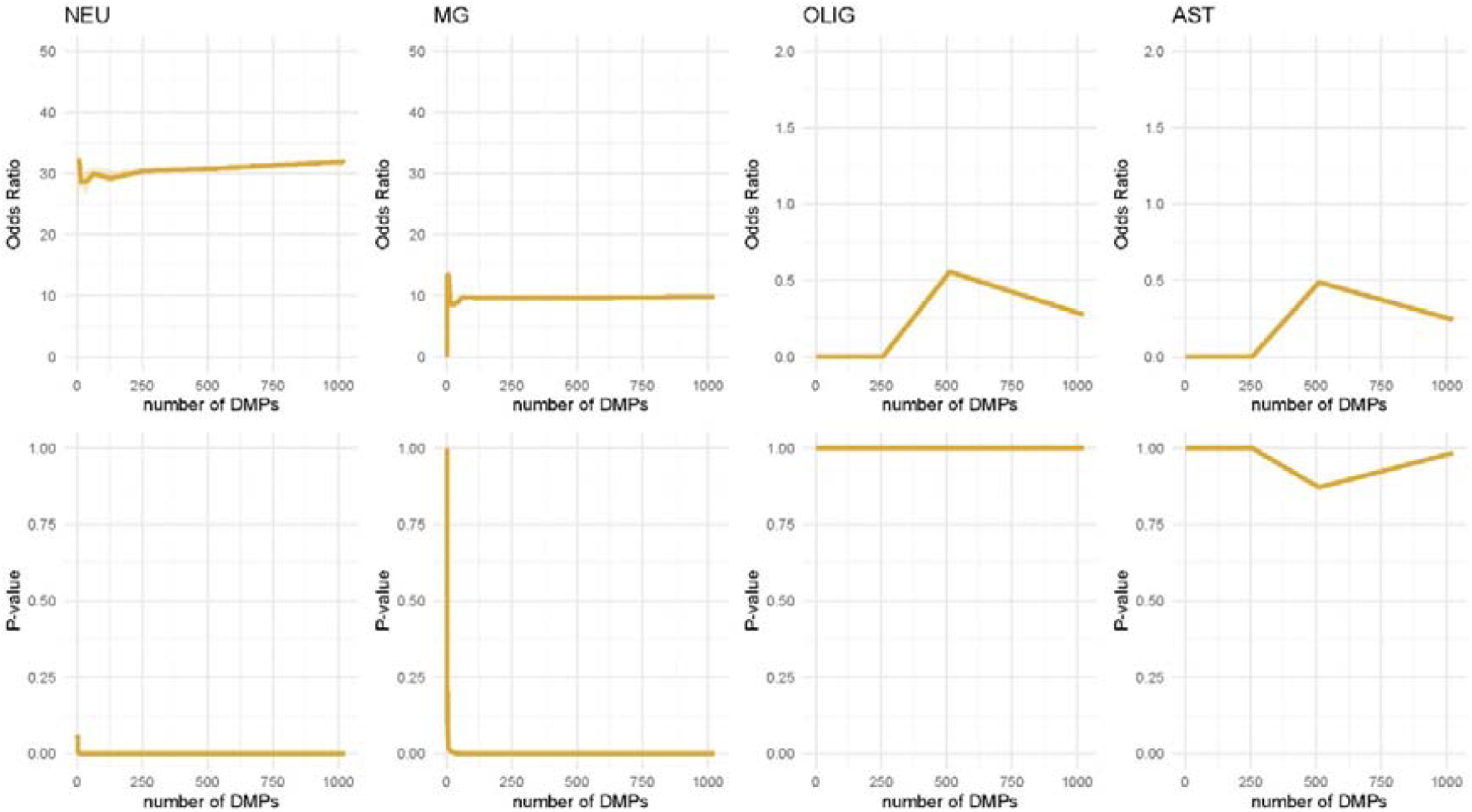
Validation on simulated data with designated DMPs in high-specificity CpG sets. Median odds ratio (OR) (upper panel) and p-value (lower panel) (Y-axis) over scenarios of increasing DMPs added to the data (X-axis). Each median was computed over 10 iterations and the ribbons show the interquartile range. From left to right, the enrichment results in each simulated cell type-specific CpG set for neurons (NEU), microglia (MG), oligodendrocytes (OLIG) and astrocytes (AST). Both neurons and microglia are found to be enriched across the majority of number of DMPs introduced in the data, which is in line with the number of DMPs assigned to their set-specific CpGs (50% to neurons and 20% to microglia). Similarly, the cell types with fewest DMPs assigned (0.1% each), oligodendrocytes and astrocytes, are not found as enriched across any amount of DMPs introduced to the data.

### Application to Brain EWAS findings

The ORA-based cell type enrichment approach was next evaluated for its ability to detect known cell type enrichment patterns in several NDDs. For this purpose, published EWASs or summary statistics were obtained for AD, LBD, and MS, and the identified DMPs were used as input for the tool. While applying all three specificity levels to the same dataset would not normally be required when using this tool in the future, this comprehensive approach was used here to systematically assess tool performance.

### AD-associated DMPs were enriched in glia

A set of 236 DMPs associated with Braak NFT stage was selected from a large meta-analysis of AD EWAS in the prefrontal cortex [2]. Cell type enrichment analysis using high-specificity CpG sets revealed significant enrichment exclusively in microglia **(Table 1 and Supplementary Table 1)**. In the medium- and low-specificity sets, microglia remained significantly enriched, with astrocyte enrichment also detected, primarily through CpGs shared between astrocytes and microglia (**Supplementary Figure 6**).

**Table 1:**
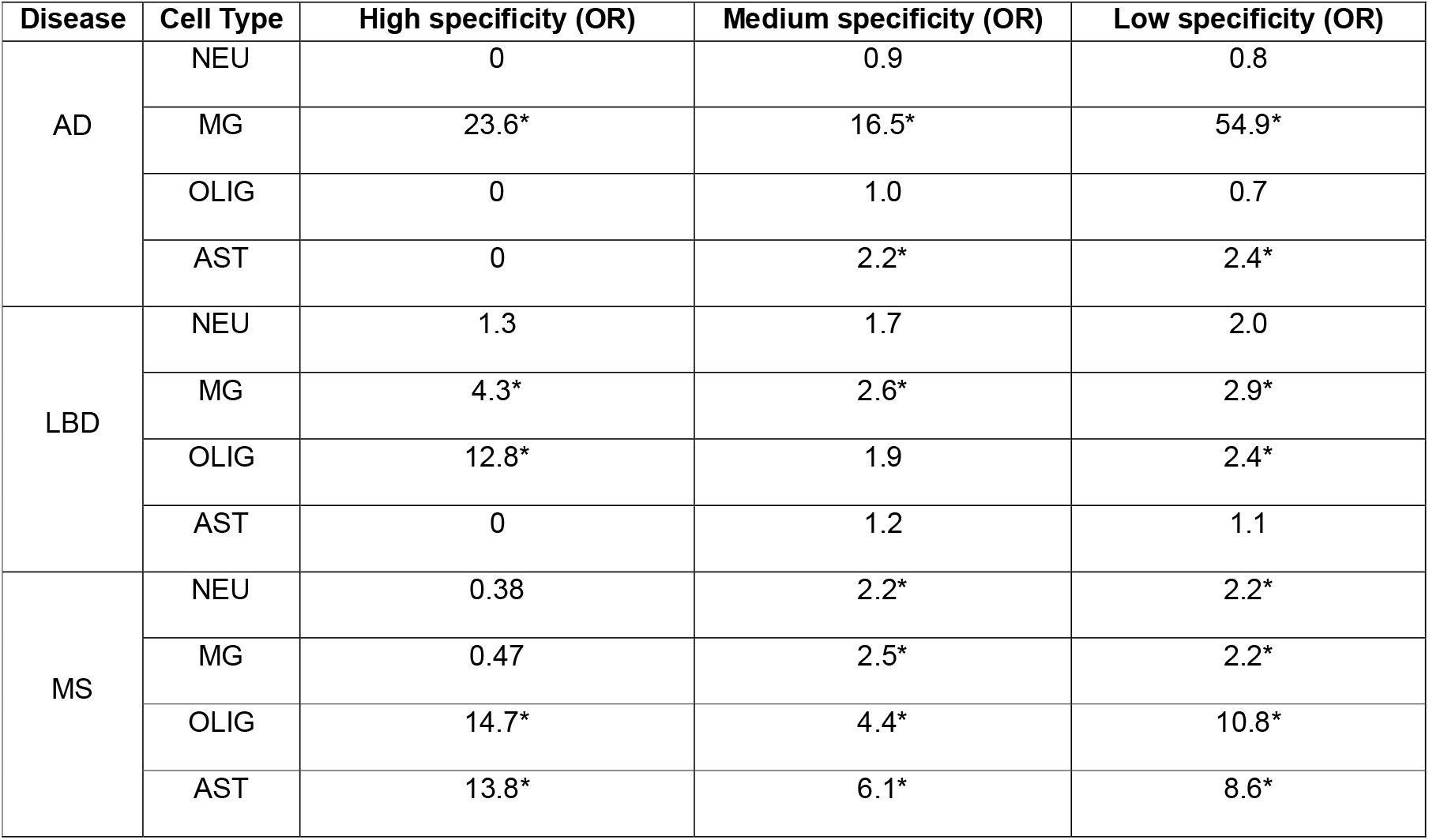
Application of the different specificity cell-type CpG sets to NDD EWAS results. The cell type enrichment tool was applied to EWASs previously undertaken in AD [2], LBD [20] and MS white matter [19]. Shown is the odds ratio (OR) for enrichment in the different cell types using the high-, medium- and low-specificity CpG sets. * Indicates significant enrichment with p-value < 0.05.

### LBD-associated DMPs were enriched in microglia and oligodendrocytes

The enrichment approach was also applied to results from an EWAS meta-analysis of Braak LB stage conducted in post-mortem pre-frontal cortex and cingulate gyrus tissue [20] using the 72 DMPs identified in that study at a suggestive p-value threshold (P < 1×10?L) out of 774,310 tested CpGs. High-specificity enrichment revealed significant overrepresentation in microglia and oligodendrocytes **(Table 1 and Supplementary Table 2**). With medium-specificity sets, only microglia showed significant enrichment, whereas both microglia and oligodendrocytes were enriched in the low-specificity sets. In all cases, enrichment was primarily attributable to CpGs not shared across cell types (**Supplementary Figure 7**).

### MS white matter-associated DMPs were enriched in oligodendrocytes

Finally, the tool was applied to results from an EWAS of MS post-mortem white matter tissue [19], serving as a control as microglia and neurons are not present in this tissue context and therefore no enrichment in these cell types should be observed. Indeed, high-specificity analysis of the 8,336 DMPs reported in the MS white-matter EWAS revealed significant enrichment in oligodendrocytes and astrocytes. In contrast, both medium- and low-specificity analyses showed significant enrichment across all cell types; however, the highest ORs were observed for oligodendrocytes and astrocytes (**Table 1 and Supplementary Table 3**).

### CEAM Web Application

To facilitate widespread adoption and ensure accessibility for the research community, we implemented CEAM as a user-friendly interactive web application using R Shiny (https://um-dementia-systems-biology.shinyapps.io/CEAM/). The app allows users to upload their own EWAS summary statistics, select reference CpG panels based on specificity, and perform cell type enrichment analysis directly through the web interface. All computations are handled on the server side, requiring no specialized programming skills.

Users receive results including odds ratios, significance, and interactive plots summarizing cell type enrichment. Outputs can be downloaded for further analysis or integration into downstream workflows. The app also provides detailed documentation, guidance on data formatting, and example datasets for new users.

We validated the CEAM app by analyzing both simulated datasets and publicly available EWAS results from neurodegenerative disease studies. In each case, the web app replicated the findings reported in this manuscript, providing consistent enrichment patterns and robust visual outputs (see **Table 1** and **Supplementary Figures 6-7**). Iterative testing and user feedback guided the design, ensuring intuitive experience and clear workflow navigation.

## Discussion

This study introduces CEAM as a robust and interpretable framework for cell type enrichment analysis of bulk brain EWAS findings. The approach adapts the ORA using cell type-specific CpG sets constructed from purified cell DNA methylation data. By systematically evaluating the approach in both simulated data and real EWAS, we demonstrate that our method is not only flexible and broadly applicable but also yields biologically meaningful results.

Through simulation experiments, we established the method’s reliability under shifts in cell type composition between sample groups as common confounding scenarios in EWASs. This is particularly important in studies of brain disorders, where shifts in the relative abundance of neurons, microglia, and other cell types can occur both as pathological hallmarks of disease progression and as artifacts introduced during tissue handling procedures. We also successfully demonstrated that accounting for cell type composition reliably prevents false-positive cell-type enrichment results, while neglecting this step leads to spurious findings. Importantly, the method showed robustness across a broad range of input sizes, accurately identifying cell type enrichment patterns even with very few DMPs. This flexibility ensures applicability across studies varying in power or effect size, making it equally valuable for large-scale meta-analyses and focused investigations.

Applying our approach to real EWAS findings from AD, LBD, and MS provided strong evidence for its biological relevance. Cell type enrichment in AD demonstrated strong microglial and astrocyte enrichment, concordant with the non-neuronal signal found in one of the replication cohorts of the original study [3]. Moreover, microglia have a pivotal role in AD onset and progression through microglial activation, which can be caused through DNAm alterations [21-23]. Enrichment results in LBD align well with previous findings, including studies reporting enrichment of DMPs associated with synaptic dysfunction and inflammation-related gene ontology terms [20]. Additionally, differential gene expression analyses have identified significant enrichment of pathways involved in oligodendrocyte development and myelination[24]. Together, these results suggest that methylation alterations observed in LBD may primarily affect microglia and oligodendrocytes, consistent with the current cell type enrichment findings. Indeed, changes in microglial activity have previously been linked specifically to both early and advanced stages of LBD pathology [25,26]. Lastly, applying the high-specificity CpG sets to the MS white matter EWAS correctly resulted in enrichment of cell types known to be abundant in white matter, demonstrating the strong discriminatory power of these sets. However, the medium- and low-specificity sets yielded less precise enrichment results, indicating a potential limitation of the tool when broader specificity criteria are used.

CEAM offers clear and straightforward interpretability, overcoming many of the limitations associated with existing deconvolution-based enrichment methods. Its consistent performance across multiple neurodegenerative diseases, each characterized by distinct cellular pathologies, underscores the versatility and broad applicability of this approach to various brain-related disorders. A key advantage of CEAM is its accessibility as a web-based tool, enabling researchers without programming expertise to easily perform cell type enrichment analysis on their own EWAS results. As new reference datasets become available, CEAM can be readily updated to incorporate additional tissues or cell types, further broadening its utility to the wider research community.

Several important limitations should be considered regarding the current methodology. First, the specificity of oligodendrocyte and astrocyte panels may be reduced due to overlapping intermediate methylation profiles, possibly resulting from technical factors such as incomplete marker resolution during nuclei sorting. Additionally, the astrocyte fraction is defined by exclusion criteria (NeuN–/SOX10–/IRF8–), inherently including other minority glial or non-neuronal populations. Thus, future refinements in sorting protocols or incorporating additional markers could enhance cell type resolution. Second, the current approach defines cell type-specific CpGs based on intermediate methylation values using empirical thresholds. Integrating DNAm data with chromatin accessibility and transcriptional profiles or employing ranking-based methods to capture a gradient of cell type association may further improve CpG selection. Finally, extending reference panels to include additional cell populations, specific disease states, or developmental stages would significantly enhance the resolution and applicability of this enrichment method.

In summary, this study establishes a validated, interpretable, and adaptable approach for cell type enrichment analysis in DNAm data, offering robust performance across common EWAS challenges and enabling direct biological interpretation. The open framework, flexibility across input sizes, and applicability to diverse NDDs position this method as a valuable addition to the epigenetics toolbox. With further refinement and expanded reference data, the approach has strong potential for broader use in both brain and non-brain tissues, empowering more nuanced, cell type-resolved insights from bulk DNAm studies.

## References

[1] Jirtle RL, Skinner MK. Environmental epigenomics and disease susceptibility. Nat Rev Genet. 2007;8(4):253–262.

[2] Smith RG, Pishva E, Shireby G, et al. A meta-analysis of epigenome-wide association studies in Alzheimer’s disease highlights novel differentially methylated loci across cortex. Nat Commun. 2021;12:3517.

[3] Shireby G, Dempster EL, Policicchio S, et al. DNA methylation signatures of Alzheimer’s disease neuropathology in the cortex are primarily driven by variation in non-neuronal cell-types. Nat Commun. 2022;13(1):5620.

[4] Zhang L, Silva TC, Young JI, et al. Epigenome-wide meta-analysis of DNA methylation differences in prefrontal cortex implicates the immune processes in Alzheimer’s disease. Nat Commun. 2020;11(1):6114.

[5] Gu JF, Barrera J, Yun Y, et al. Cell-Type Specific Changes in DNA Methylation of Intron 1 in Synucleinopathy Brains. Front Neurosci-Switz. 2021;15.

[6] De Ridder K, Che HW, Leroy K, Thienpont B. Benchmarking of methods for DNA methylome deconvolution. Nat Commun. 2024;15(1).

[7] Skene NG, Grant SGN. Identification of Vulnerable Cell Types in Major Brain Disorders Using Single Cell Transcriptomes and Expression Weighted Cell Type Enrichment. Front Neurosci-Switz. 2016;10.

[8] Breeze CE, Paul DS, van Dongen J, et al. eFORGE: A Tool for Identifying Cell Type-Specific Signal in Epigenomic Data. Cell Rep. 2016;17(8):2137–2150.

[9] Zheng SJC, Breeze CE, Beck S, Teschendorff AE. Identification of differentially methylated cell types in epigenome-wide association studies. Nat Methods. 2018;15(12):1059-+.

[10] Francis PT, Costello H, Hayes GM. Brains for Dementia Research: Evolution in a Longitudinal Brain Donation Cohort to Maximize Current and Future Value. J Alzheimers Dis. 2018;66(4):1635–1644.

[11] Shireby G, Dempster EL, Policicchio S, et al. DNA methylation signatures of Alzheimer’s disease neuropathology in the cortex are primarily driven by variation in non-neuronal cell-types. Nat Commun. 2022;13(1).

[12] Pidsley R, Wong CCY, Volta M, et al. A data-driven approach to preprocessing Illumina 450K methylation array data. Bmc Genomics. 2013;14.

[13] Triche TJ, Weisenberger DJ, Van Den Berg D, et al. Low-level processing of Illumina Infinium DNA Methylation BeadArrays. Nucleic Acids Res. 2013;41(7).

[14] Teschendorff AE, Marabita F, Lechner M, et al. A beta-mixture quantile normalization method for correcting probe design bias in Illumina Infinium 450 k DNA methylation data. Bioinformatics. 2013;29(2):189–196.

[15] McCartney DL, Walker RM, Morris SW, et al. Identification of polymorphic and off-target probe binding sites on the Illumina Infinium MethylationEPIC BeadChip. Genom Data. 2016;9:22–4.

[16] Leek JT, Johnson WE, Parker HS, et al. sva: Surrogate variable analysis. 2024.

[17] Hannon E, Knox O, Sugden K, et al. Characterizing genetic and environmental influences on variable DNA methylation using monozygotic and dizygotic twins. PLoS Genet. 2018;14(8):e1007544.

[18] Weber F, Knapp G, Ickstadt K, et al. Zero-cell corrections in random-effects meta-analyses. Research Synthesis Methods. 2020;11(6):913–919.

[19] Tiane A, Schepers M, Reijnders RA, et al. From methylation to myelination: epigenomic and transcriptomic profiling of chronic inactive demyelinated multiple sclerosis lesions. Acta Neuropathologica. 2023;146(2):283–299.

[20] Harvey J, Imm J, Kouhsar M, et al. Interrogating DNA methylation associated with Lewy body pathology in a cross brain-region and multi-cohort study. medRxiv. 2025:2025.03.13.25323837.

[21] Miao J, Ma H, Yang Y, et al. Microglia in Alzheimer’s disease: pathogenesis, mechanisms, and therapeutic potentials. Frontiers in Aging Neuroscience. 2023;15:1201982.

[22] Dias D, Socodato R. Beyond Amyloid and Tau: The Critical Role of Microglia in Alzheimer’s Disease Therapeutics. Biomedicines. 2025;13(2):279.

[23] Wang L, Yu C-C, Liu X-Y, et al. Epigenetic Modulation of Microglia Function and Phenotypes in Neurodegenerative Diseases. Neural Plast. 2021;2021:9912686.

[24] Pietrzak M, Papp A, Curtis A, et al. Gene expression profiling of brain samples from patients with Lewy body dementia. Biochem Biophys Res Commun. 2016;479(4):875–880.

[25] Surendranathan A, Su L, Mak E, et al. Early microglial activation and peripheral inflammation in dementia with Lewy bodies. Brain. 2018;141(12):3415–3427.

[26] Streit WJ, Xue Q-S. Microglia in dementia with Lewy bodies. Brain, Behavior, and Immunity. 2016;55:191–201.

