## Supplementary material for "A Cell Type Enrichment Analysis Tool for Brain DNA Methylation Data (CEAM)": Supplgurementary Figures

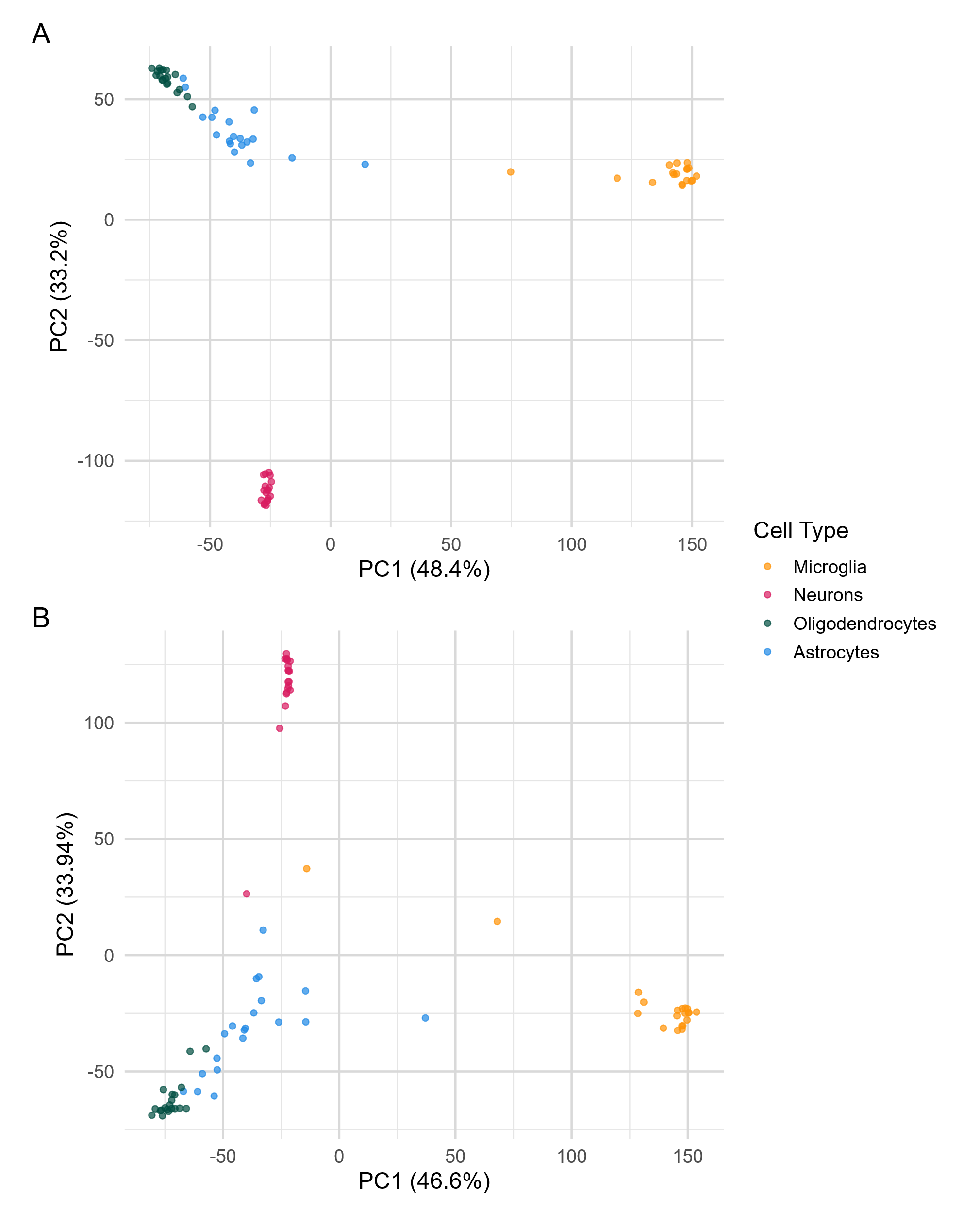

**Supplementary Figure 1. Visualization of the first two principal components of DNAm beta-values in four cell types, across control samples from two independent cohorts.** (A) UKBBN cohort: samples cluster distinctly by cell type, with minor overlap between astrocytes and oligodendrocytes. (B) BDR cohort: similar cell type separation is observed. Plots were examined to assess cell type grouping and identify potential sources of confounding variation.

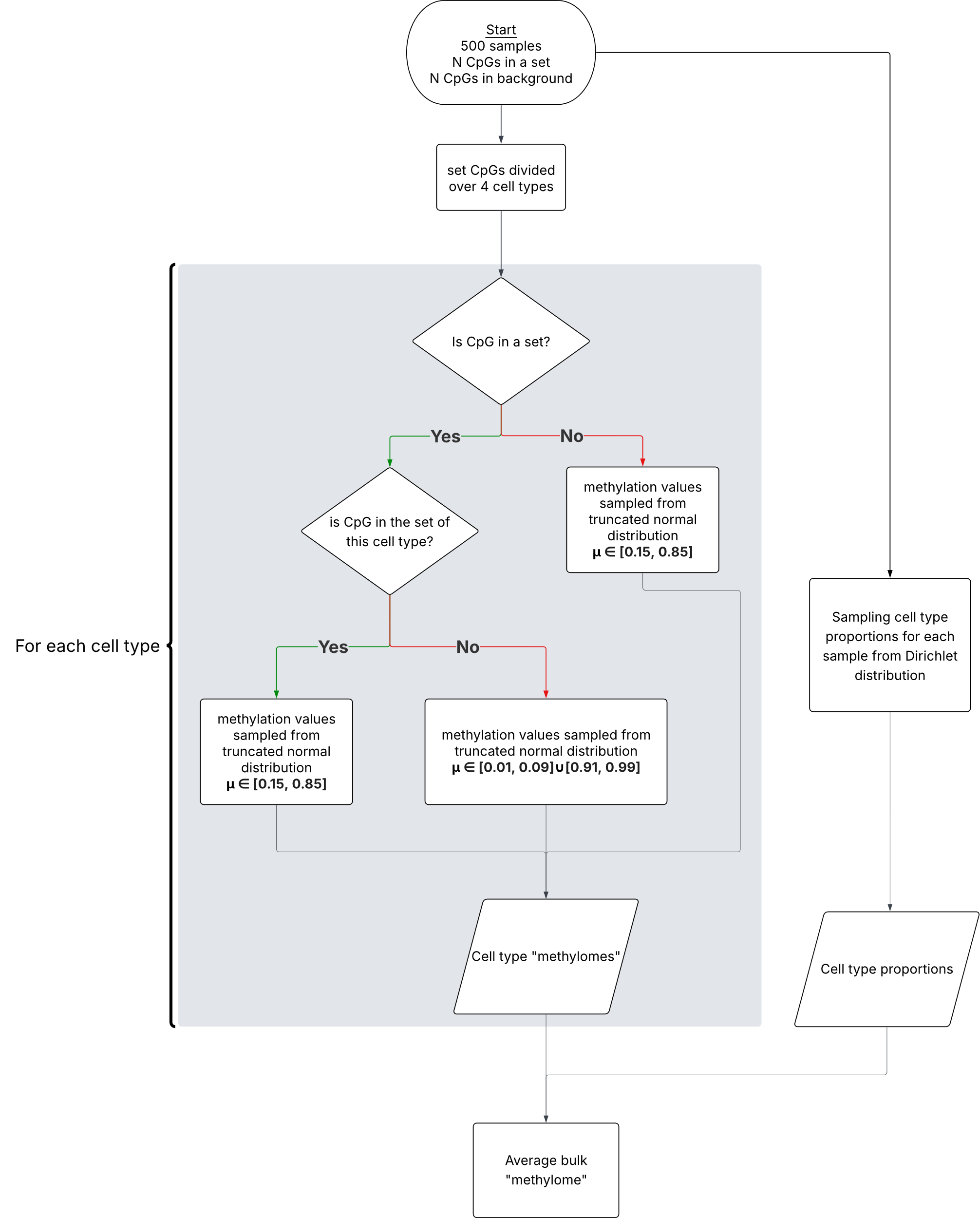

**Supplementary Figure 2. Flowchart demonstrating how DNAm data was simulated.** Methylation values were assigned to each CpG for each cell type and subsequently summed up according to simulated cell type proportions to generate “pseudobulk” samples.

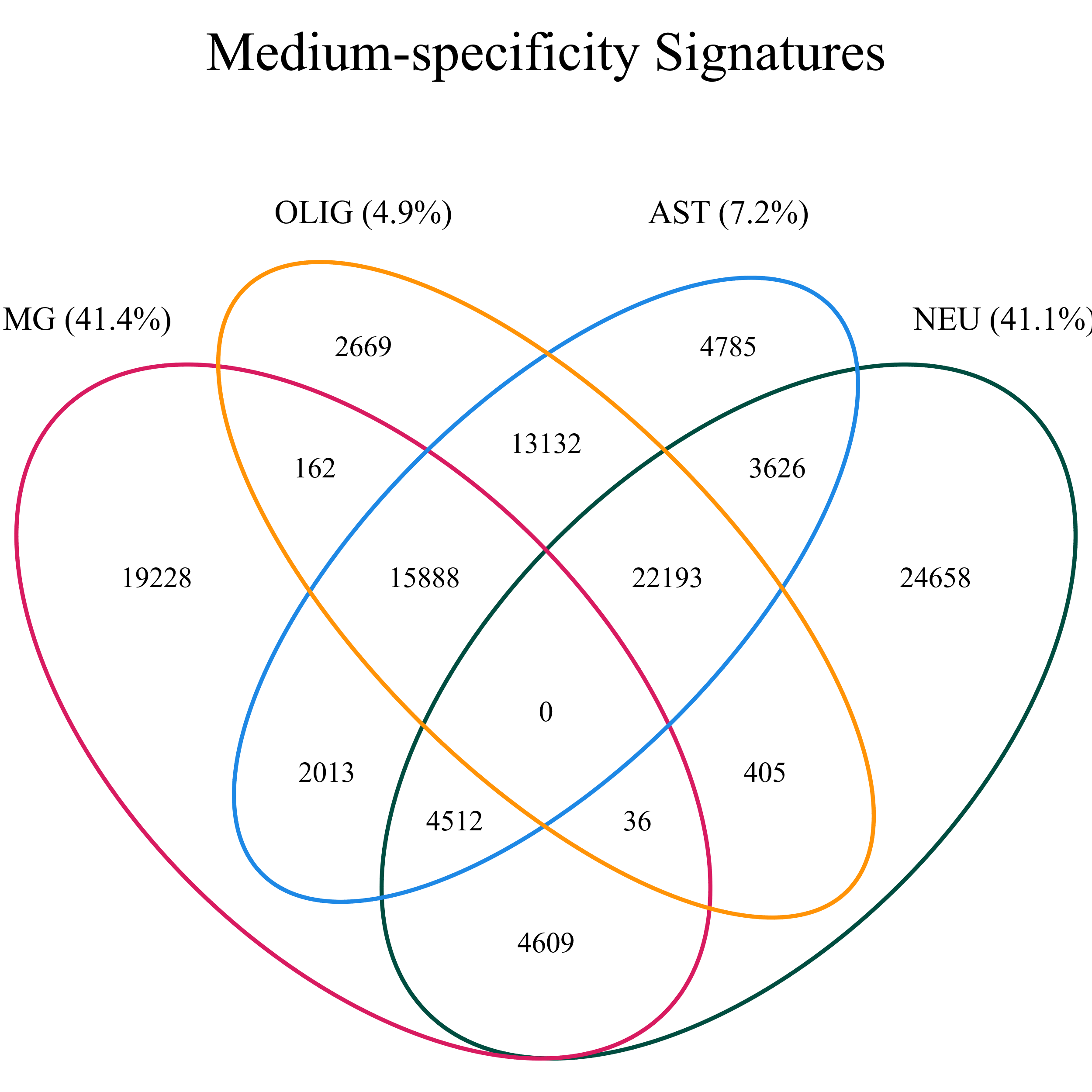

**Supplementary Figure 3. Composition of medium-specificty CpG sets.** The venn diagram shows how many CpGs are unique for each CpG set (indicated by the %) and how many CpGs overlap between sets for microglia (MG), oligodendrocytes (OLIG), astrocytes (AST) and neurons (NEU).

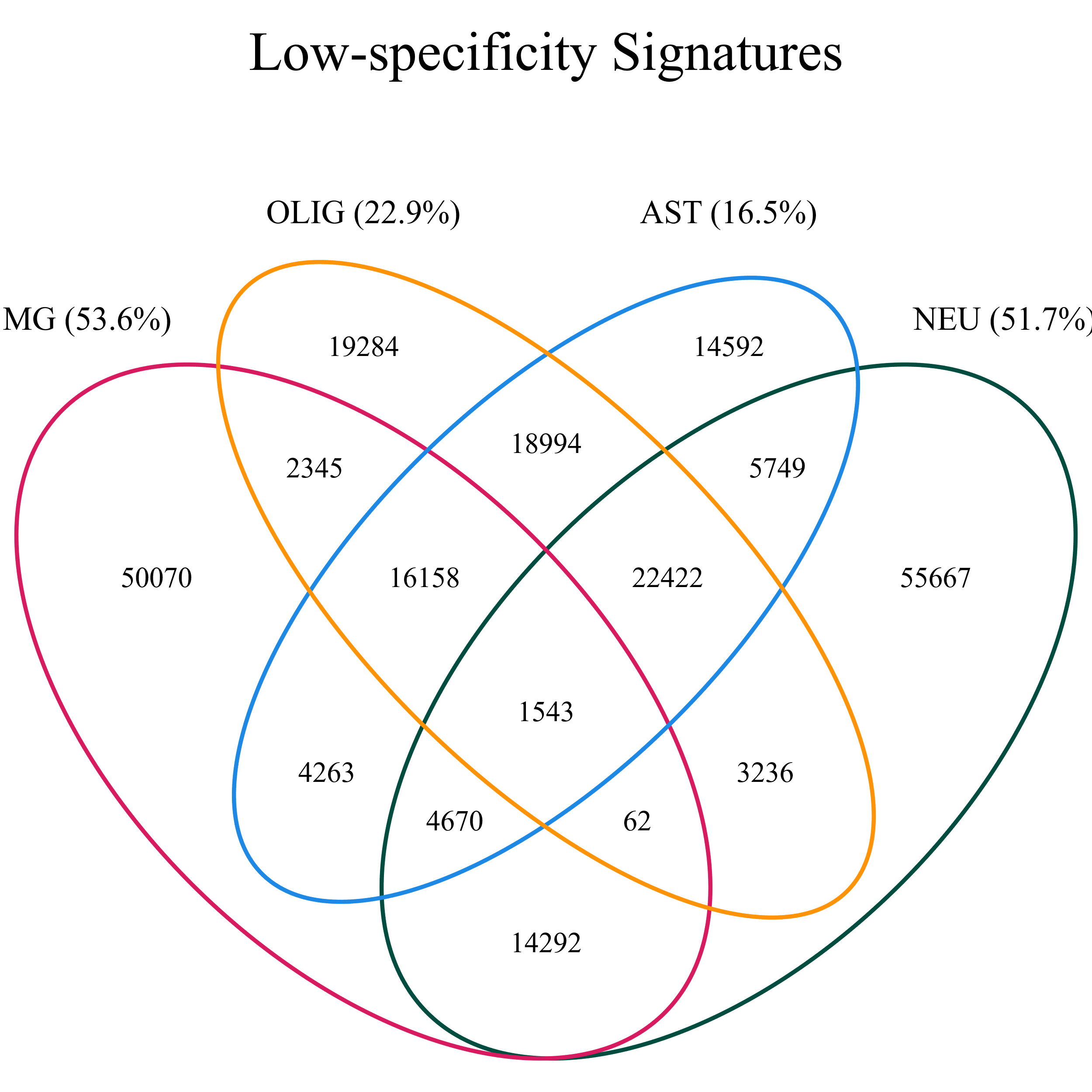

**Supplementary Figure 4. Composition of low-specificty CpG sets.** The venn diagram shows how many CpGs are unique for each CpG set (indicated by the %) and how many CpGs overlap between sets. Notably the number of unique CpGs increased for all cell types compared to the medium-specificity sets for microglia (MG), oligodendrocytes (OLIG), astrocytes (AST) and neurons (NEU).

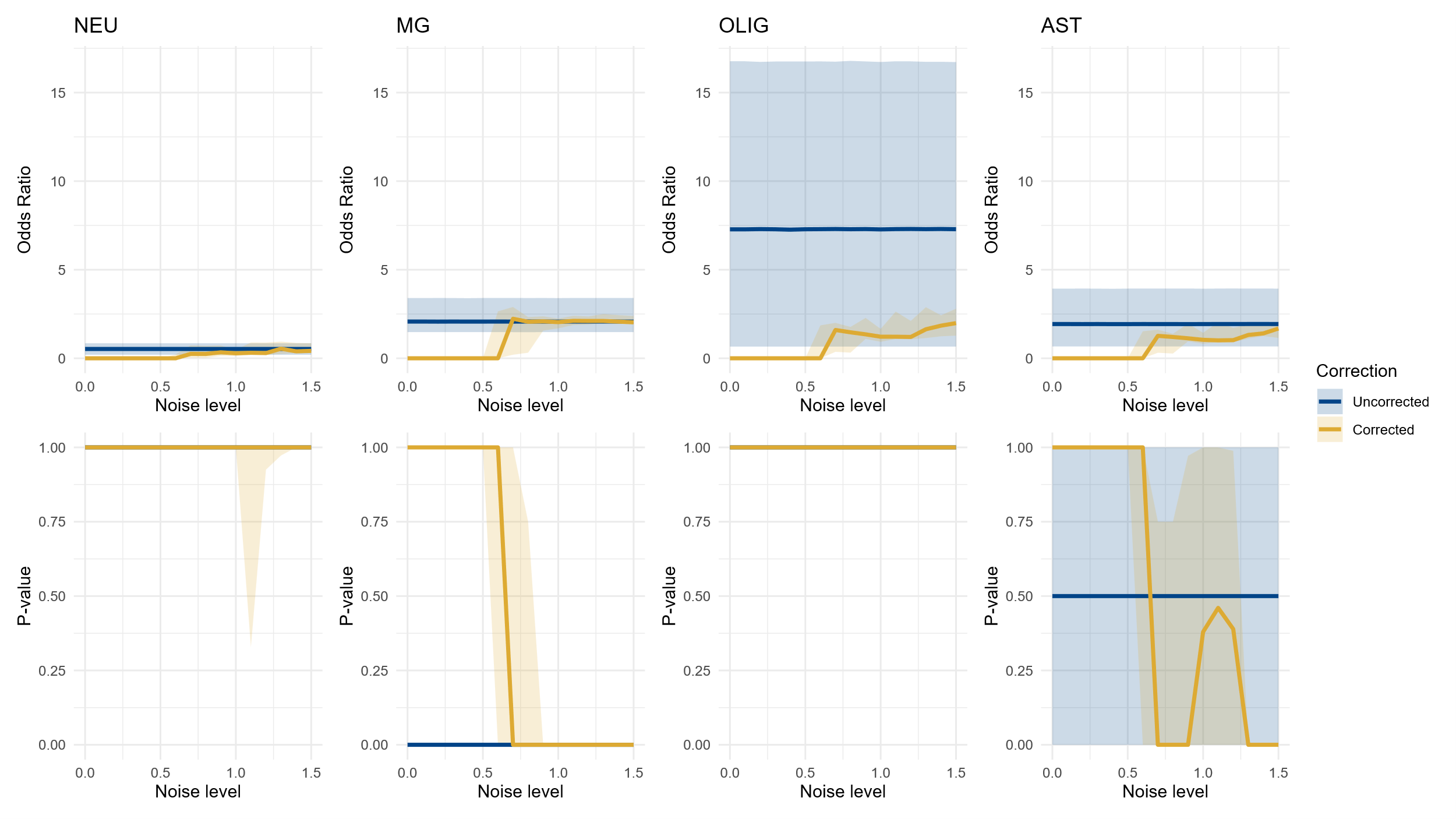

**Supplementary Figure 5. Validation on simulated data with shifted cell type proportions in simulated medium-specificity CpG sets.** Median odds ratio (OR) (upper panel) and P-value (lower panel) (Y-axis) over scenarios of increasing noise in cell type proportions used for correction (X-axis). Each median was computed over 10 iterations and the ribbons show the interquartile range (IQR). From left to right, the enrichment results in each simulated cell type-specific CpG set for neurons (NEU), microglia (MG), oligodendrocytes (OLIG) and astrocytes (AST), once uncorrected for cell type composition (yellow) and once corrected (blue). A relative reduction of neurons in the simulated data causes microglia and astrocytes to become significantly enriched in the uncorrected scenario with P-values < 0.05. However, correcting the analysis with known cell type proportions returns no enrichment in any cell type. After adding enough noise (noise level = 0.5) microglia and astrocytes become similarly enriched to the uncorrected results. In instances where not a single CpG of a set was in the input and therefore no OR could be computed, the OR was set to 0 for visualization purposes.

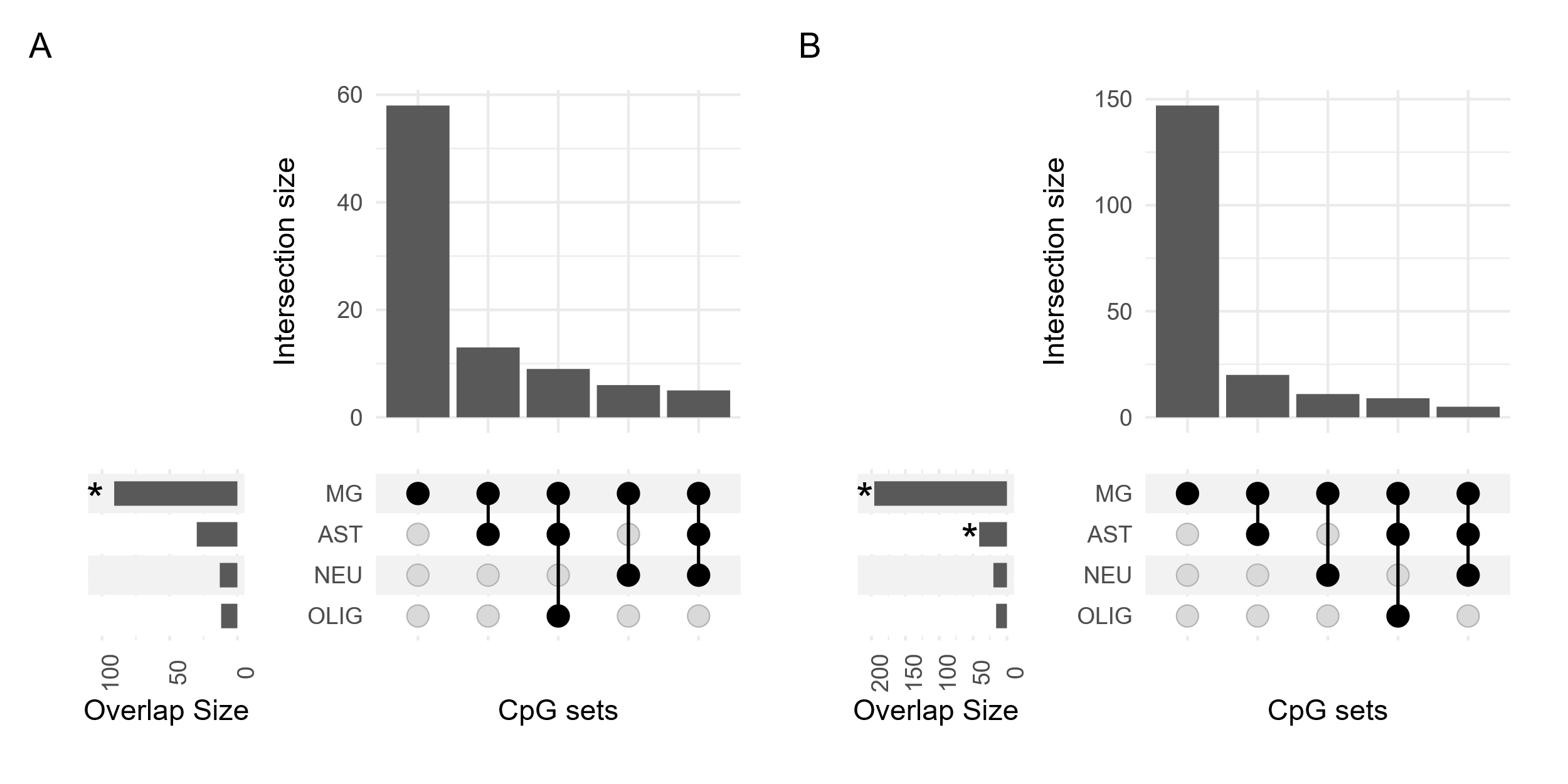

**Supplementary Figure 6. Upset plot visualization of cell type enrichment in 236 AD-associated DMPs.** (A) Using medium-specificity CpG sets, the AD-associated DMPs were found to be enriched in microglia, with predominantly CpGs unique to microglia driving the enrichment. (B) Cell type enrichment using low-specificity CpG sets yielded similar enrichment patterns, although in this instance astrocytes now showed enrichment, although this was driven through shared CpGs with microglia. * Indicates false discovery rate (FDR) significance, defined as q-value < 0.05. Cell types are abbreviated as follows: MG = microglia, AST = astrocytes, NEU = neurons, and OLIG = oligodendrocytes.

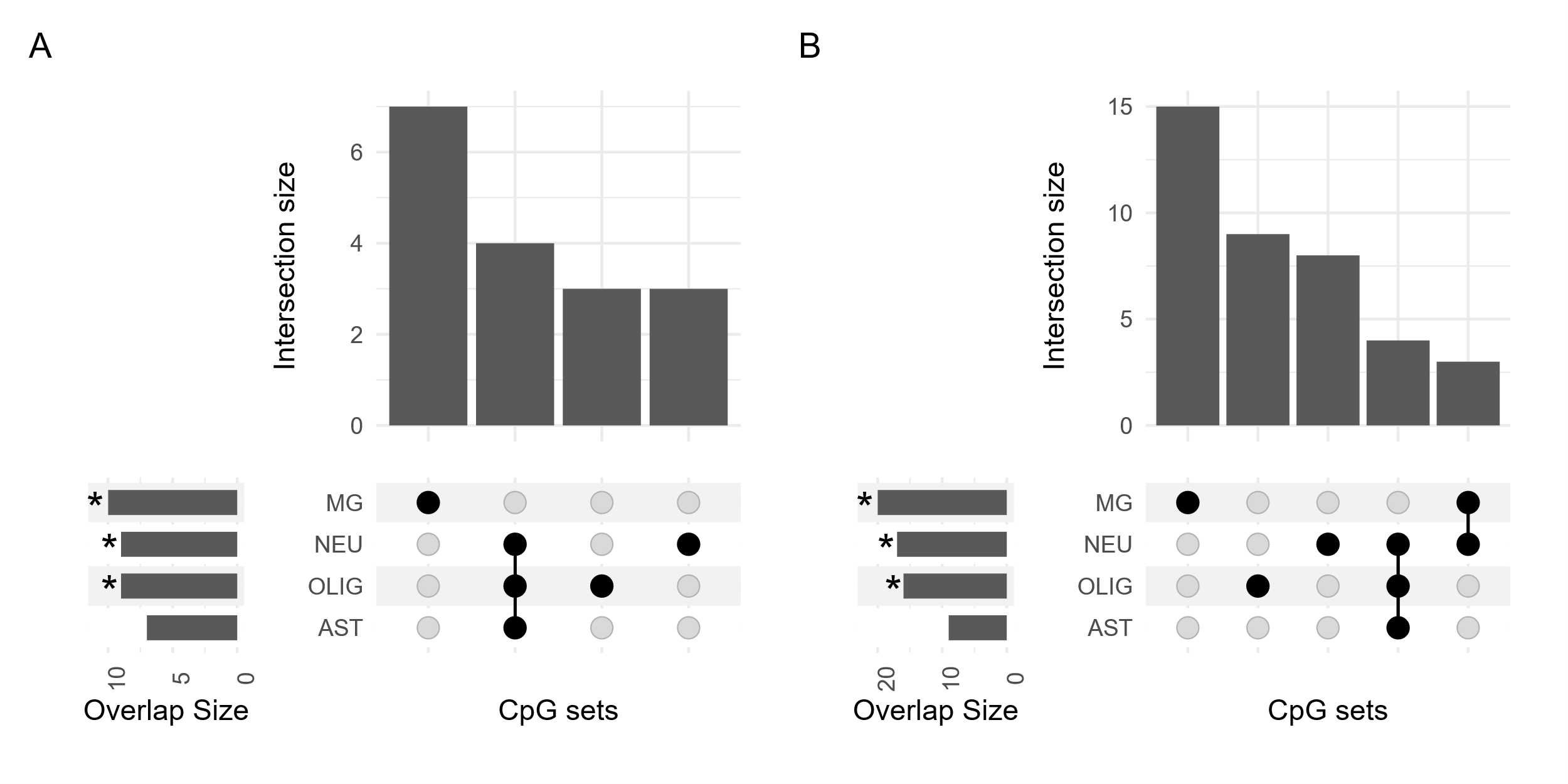

**Supplementary Figure 7. Upset plot visualization of cell type enrichment in 71 LBD-associated CpGs.** (A) When using the medium-specificity CpG sets microglia, neurons and oligodendrocytes are enriched. (B) The same enrichment patterns persist when using low-specificity CpG sets. * Indicates significance as q-value < 0.05. Cell types are abbreviated as follows: MG = microglia, AST = astrocytes, NEU = neurons, and OLIG = oligodendrocytes.

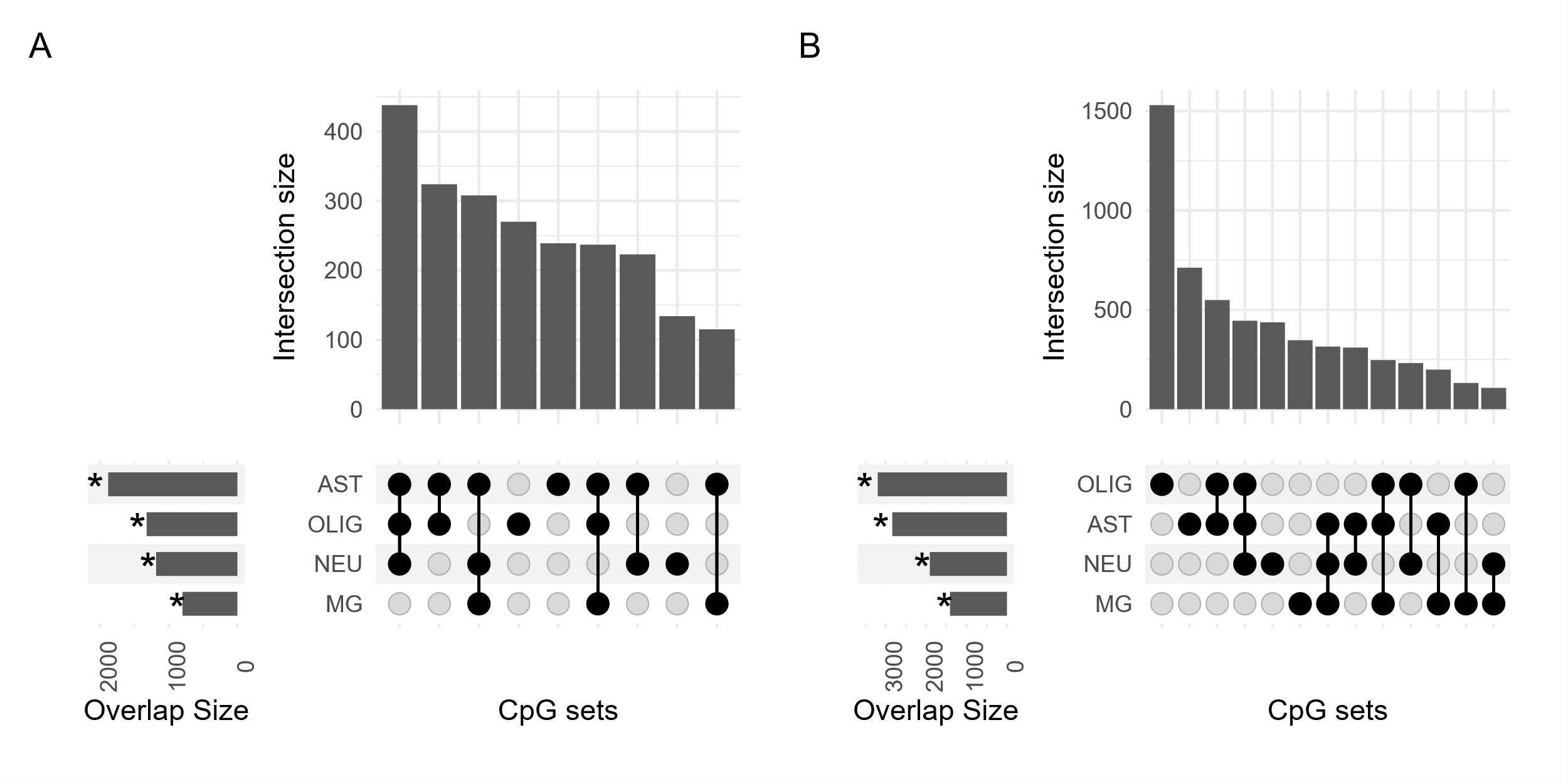

**Supplementary Figure 8. Upset plot visualization of cell type enrichment in 8336 MS-associated CpGs.** When using either the medium- (A) or low-specificity (B) CpG sets all cell types are enriched. Notably, oligodendrocytes and astrocytes remain the cell types with the largest overlap and the highest number of unique CpGs driving their enrichment.
